# Musculoskeletal architecture of the shoulder: A comparative anatomy study in bats and mice informing human rotator cuff function

**DOI:** 10.1101/2025.07.04.663211

**Authors:** Iden Kurtaliaj, Jennifer Kunes, Shicheng Li, Michael Rowley, Lynn Ann Forrester, Mikhail Golman, Sharon M. Swartz, William N. Levine, Guy M. Genin, Stavros Thomopoulos

**Author notes:** **Corresponding authors** Stavros Thomopoulos, Guy Genin.

## Abstract

Overhead motion in humans often leads to shoulder injuries, a consequence of the evolutionary trade-off in glenohumeral joint anatomy that balances stability with mobility. Bats consistently engage in overhead motion during flight, subjecting their shoulders to substantial loading throughout their relatively long lifespan. Remarkably, despite the demands placed on a bat’s shoulder, instability and rotator cuff tears, which could be fatal to bats in short order, are not observed in nature. We were thus inspired to study functional adaptations in the shoulders of bats that enable this overhead motion. Comparative anatomical studies of the shoulders of bats and mice, similarly-sized quadrupeds, were performed and interpreted using a mathematical model. Scapular anatomy indicated a more prominent role for the infraspinatus muscle in the bat compared to the mouse. Measurements of bat and mice shoulders revealed that the bat glenoid had a larger curvature and arc length than that of mice, providing a larger articulating surface area with and deeper enclosing surface of the humeral head. Modeling results predicted that the bat shoulder is stable over a dramatically larger range of angles compared to the mouse shoulder. These results suggested that adaptations to constraints imposed by the bony anatomy and rotator cuff tendons of the shoulder may contribute to the ability of bats to sustain overhead motion in a high stress, repeated loading environment without injury. Results suggest that bats have evolved unique adaptations in their glenohumeral bony anatomy that reduce stress on the supraspinatus, enhance joint stability, and optimize strength across a broad range of motion.

## Introduction

The human shoulder is a remarkable joint, allowing for an exceptional range of motion that far exceeds that of most non-primate mammals[1]. This mobility has been critical in the evolution of human behaviors such as throwing and tool use, but has come at a cost: the glenohumeral joint that enables this motion – the most mobile joint in the human body – is inherently unstable and prone to injury [2–4]. Shoulder injuries are one of the most prevalent musculoskeletal pathologies, particularly affecting athletes participating in overhead sports and workers undertaking overhead tasks. The unique biomechanical demands of such overhead activities predispose the shoulder joint to a heightened risk of so-called “overuse” pathologies [5–8]. Rotator cuff tears are one of the most common shoulder overuse injuries, affecting more 17 million individuals annually in the US, with incidence of injury increases with age to over 40% for those over 65 [9–13] and widespread impact on the healthcare system and workforce [14–18].

To accommodate large ranges of motion, the human glenohumeral joint of the human shoulder has evolved with minimal bony constraints. Unlike the hip joint, where the head of the femur is deeply embedded in the pelvis, the humeral head sits in a shallow indentation in the scapula called the glenoid fossa, a setting often compared to a golf ball on a tee [19,20]. Instability of the humeral head is mitigated through a system of muscles and tendons, collectively known as the rotator cuff, that compresses the humeral head against the glenoid fossa [14,21–24]. Nevertheless, injuries to the rotator cuff are rather common, disrupting the force balance in the glenohumeral joint and often leading to shoulder instability [5,22,24].

This is in sharp contrast to bats. Mammals of Order Chiroptera evolved wings of skin stretched between elongated fingers and a range of musculoskeletal features that enable flight. The biomechanical demands placed on a bat’s shoulder during flight are immense, estimated to exceed the cumulative stress endured by a competitive swimmer’s shoulder by 45-fold over a lifetime [25,26]. Despite these extreme demands, no evidence of shoulder overuse injuries can be found in the large literature on wild bat anatomy and physiology, even amongst bats that reach ages of 30 years or more; a search of literature concerning injuries to bats in natural conditions produces over 130 publications but no reports of shoulder morbidity among musculoskeletal trauma. Although selection bias is possible because study of wild bats involves capture during flight using nets [27,28], indirect evidence from their longevity, the lack of reported pathology, and their reliance upon flight for feeding, escape, and migration suggest that bats may have a shoulder capable of exceptional mobility and repetitive loading without the apparent instability and injury seen in human shoulders[29,30].

We therefore hypothesized that bats have evolved adaptations to shoulder anatomy that are consistent with joint stability in overhead motion. To investigate this, we compared bat shoulders to mouse shoulders through anatomical and biomechanical measurements and modeling. Mice were used because they share aspects of shoulder anatomy with humans, including having the supraspinatus tendon pass below the coracoacromial arch, a feature rarely observed in other mammals and are widely used as an animal model for pathological conditions of the human shoulder [31,32]. Bats also share similar coracoacromial arch anatomy with humans, but use overhead motion for flight, live ten times longer than mice, and experience approximately 30-fold greater shoulder loading each day than mice [33,34]. By quantifying the anatomical features and biomechanical properties of these two similarly sized mammals, we identified several distinctive adaptations that could potentially inform new approaches for treating and preventing human shoulder pathology.

## Results

### Scapular anatomy suggests a more prominent role for the infraspinatus muscle in the bat compared to the mouse

We compared the scapulae of *C. perspicillata* and quadrupedal mice and determined linear and areal supraspinatus and infraspinatus indices. These indices describe the proportions of the scapula and provide insights into the function and mobility-stability trade-offs of the shoulder joint. Anatomical measurements showed that the scapular and infraspinatus linear indices were significantly larger in *C. perspicillata* relative to mice, whereas the supraspinatus index did not differ between the two species (Figure 2A). Similarly, the infraspinatus area index was significantly larger whereas the supraspinatus area index was significantly smaller in *C. perspicillata* relative to mice (Figure 2B).

**Figure 1.**
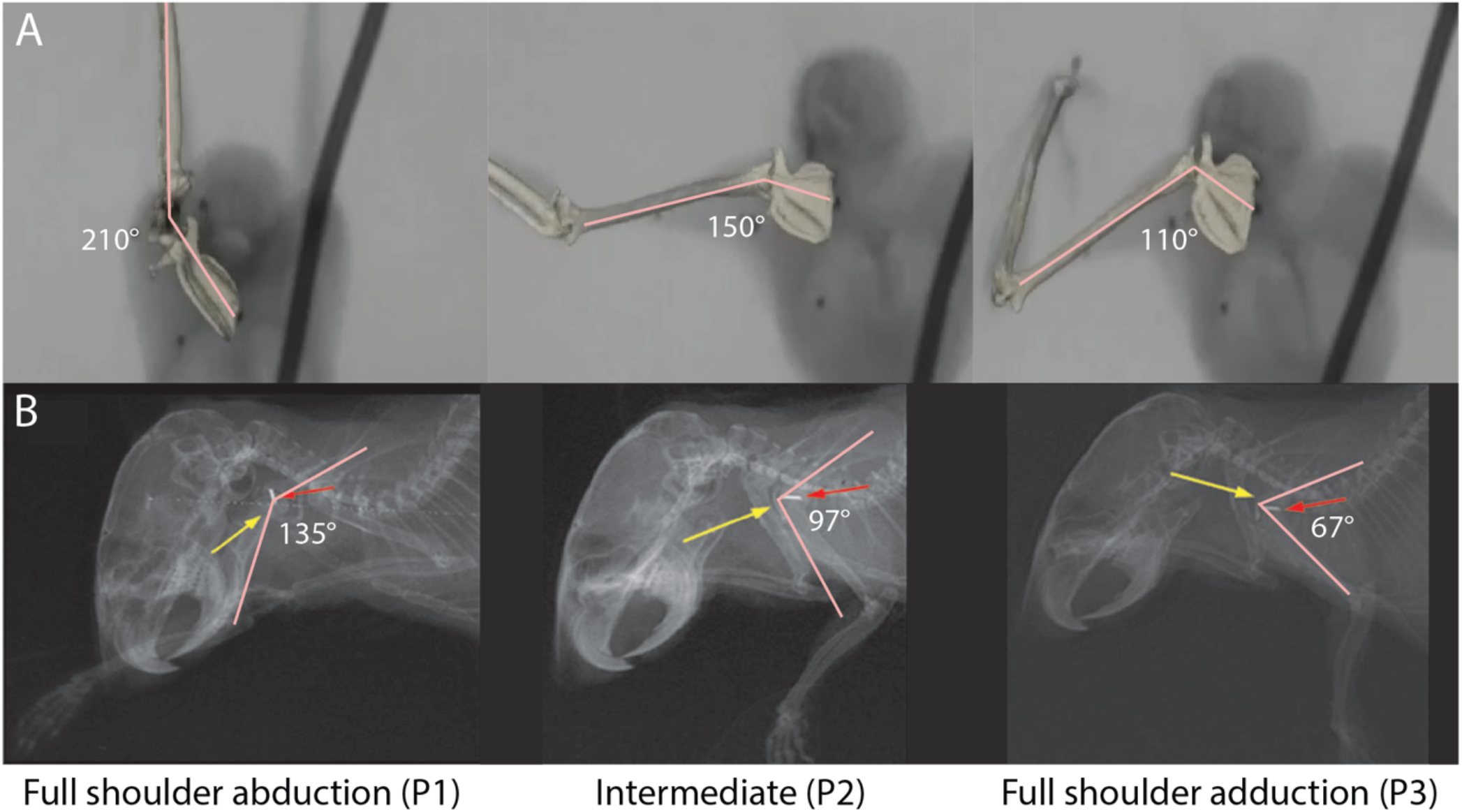
Angle between the humerus and spine of the scapula of *C. perspicillata* (A) and mice (B). Glenohumeral joints, rotator cuff muscles, and other surrounding tissue were dissected and then fixed in three different positions: full shoulder extension (“P1”), intermediate position (“P2”), and full shoulder flexion (“P3”). To consistently identify the angle between the scapular spine and the humerus for each fixation position, gait (mouse) and flight (bat) analyses were used. Adapted, with permission, from Soslowsky et al. *[31]* and from Konow et al. *[68]*.

**Figure 2.**
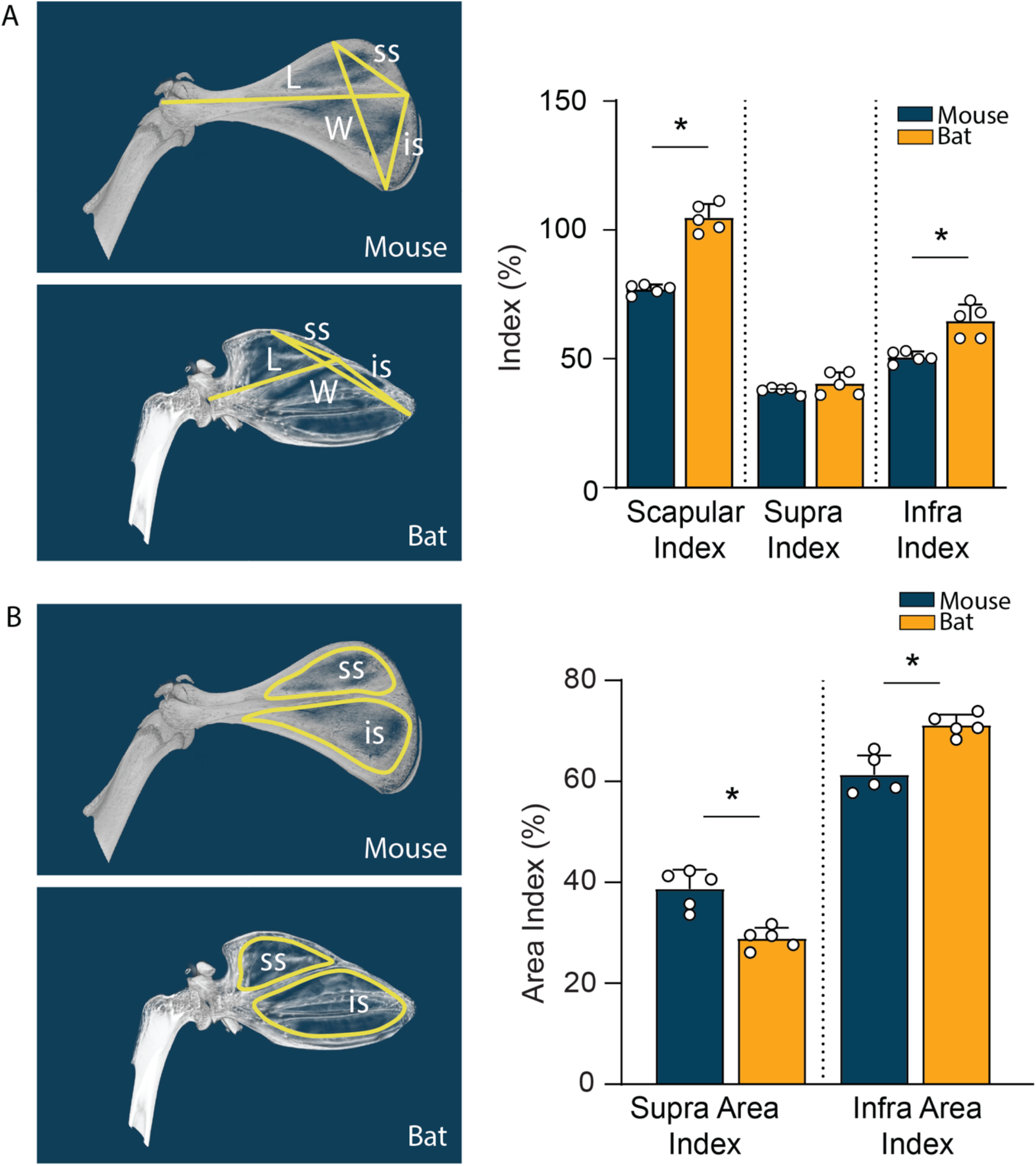
(A) Scapular, supraspinatus and infraspinatus indices. The scapular and infraspinatus indices were significantly larger in bats than in mice. There was no significant difference in the supraspinatus index between species. **(B) Supraspinatus and infraspinatus area indices**. Bats showed significantly larger infraspinatus area index and significantly lower supraspinatus area index compared to mice.

### Enlarged supraspinatus outlet in bats provides more space for the supraspinatus compared to the mouse

The supraspinatus outlet area is the space beneath the acromion of the scapula through which the supraspinatus tendon passes as it inserts onto the humeral head [35,36]. Clinically, its height (often referred to as “clearance”) and the magnitude of the outlet area are significant metrics, as a decrease in either can lead to impingement of the supraspinatus tendon and subsequent pathology [35–41]. If the clearance or the overall area is reduced, there is an increased risk for the supraspinatus tendon to become impinged or compressed between the humeral head and the overlying acromion, a condition referred to as subacromial impingement syndrome [35–41].

We compared the supraspinatus outlet and clearance in bats and mice across three defined shoulder positions: full shoulder extension (“P1”), intermediate posture (“P2”), and full shoulder flexion (“P3”). As the shoulder transitioned from full flexion to extension (i.e., from positions 1 through 3) there was an increase in the supraspinatus-acromion clearance for both *C. perspicillata* and mice (Figure 3C). However, *C. perspicillata* had a significantly larger supraspinatus outlet area compared to mice, suggesting that the morphology in *C. perspicillata* provides additional space beneath the coracoacromial arch for the supraspinatus tendon (Figure 3B).

**Figure 3.**
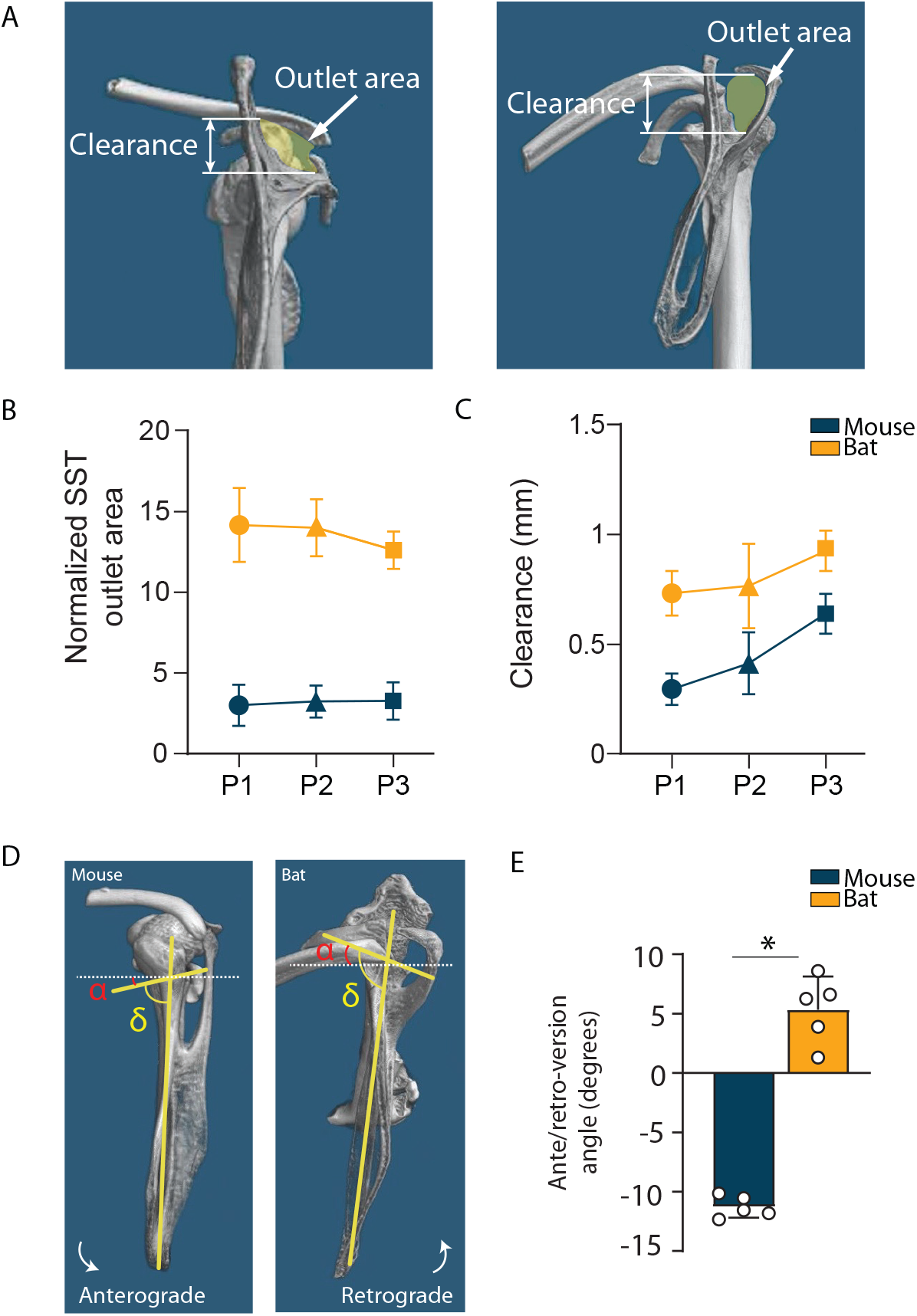
**(A)** Representative 3D microCT images of the glenohumeral joint of the *C. perspicillata* and mouse showing supraspinatus outlet area and clearance measurements. **(B) Supraspinatus outlet area (yellow in A)** normalized by supraspinatus attachment area. The normalized supraspinatus outlet area was significantly greater in *C. perspicillata* compared to mice, suggesting more space in bats for the supraspinatus to pass through to the humeral attachment. **(C) Clearance, measured as the height of the outlet area (white double-headed arrow in A)**. The supraspinatus clearances of bats and mice were measured across three defined shoulder positions:. As the shoulder transitions from full shoulder flexion to extension (i.e from full shoulder extension [“P1”] to intermediate [“P2”] to full shoulder flexion [“P3”]) there was an increase in the supraspinatus-acromion clearance for both *C. perspicillata* and mice. **(D)** Representative 3D microCT images of the glenohumeral joint of *C. perspicillata* and mouse showing glenoid version measurements. **(E)** Glenoid ante/retroversion α angle; α = δ - 90 for mice and *C. perspicillata*. The glenoid was retroverted in *C. perspicillata* and anteverted in mice.

Glenoid version describes the angular orientation between the glenoid plane and the scapular body [42]. If the glenoid plane aligns exactly with the scapular plane, it is considered neutral glenoid version; if it tilts backward (posterior), it is considered retroversion (negative version); and if it tilts forward (anterior), it is considered anteversion (positive version).This clinically relevant measure is frequently used to understand shoulder biomechanics and identify related pathologies. In humans, the typical glenoid version is close to 0°, with occasional slight anteversion or more commonly, a subtle retroversion, usually less than 10° in either direction [36,40,42–45]. Deviations from the normal version alters glenohumeral mechanics and may predispose the shoulder to instability [36,40,42–45]. Our measurements of bat and mouse shoulders revealed a retroverted glenoid in *C. perspicillata* and an anteverted glenoid in mice (Figure 3E). The retroverted orientation in *C. perspicillata* aligns with observations in high-level overhead-throwing athletes [46–48], possibly facilitating greater joint rotation while minimizing capsular tension.

### Bat glenoid anatomy contributes to greater joint stability compared to mice

Concavity compression plays a crucial role in glenohumeral stability during mid-range motion due to capsular and ligament laxity; glenoid concavity can be measured by its radius of curvature, with both radius and depth being essential for stability [20,32,49]. Measurements of bat and mice shoulders revealed that the bat glenoid had a larger curvature and arc length than that of mice, providing a larger articulating surface area that more deeply encloses the surface of the humeral head (Figure 4 C,D). In addition, the ratio of glenoid depth to humeral head width was higher in *C. perspicillata* enabling the bat glenoid to encompass a greater portion of the humeral head compared to that of the mouse (Figure 4 E,F). These findings suggest that the bat’s deeper and more encompassing glenoid contributes to enhanced bony stability in the joint. This aligns with pathological glenoids observed in patients with shoulder instability, which typically show a reduction in glenoid concavity and a more flattened morphology [50].

**Figure 4.**
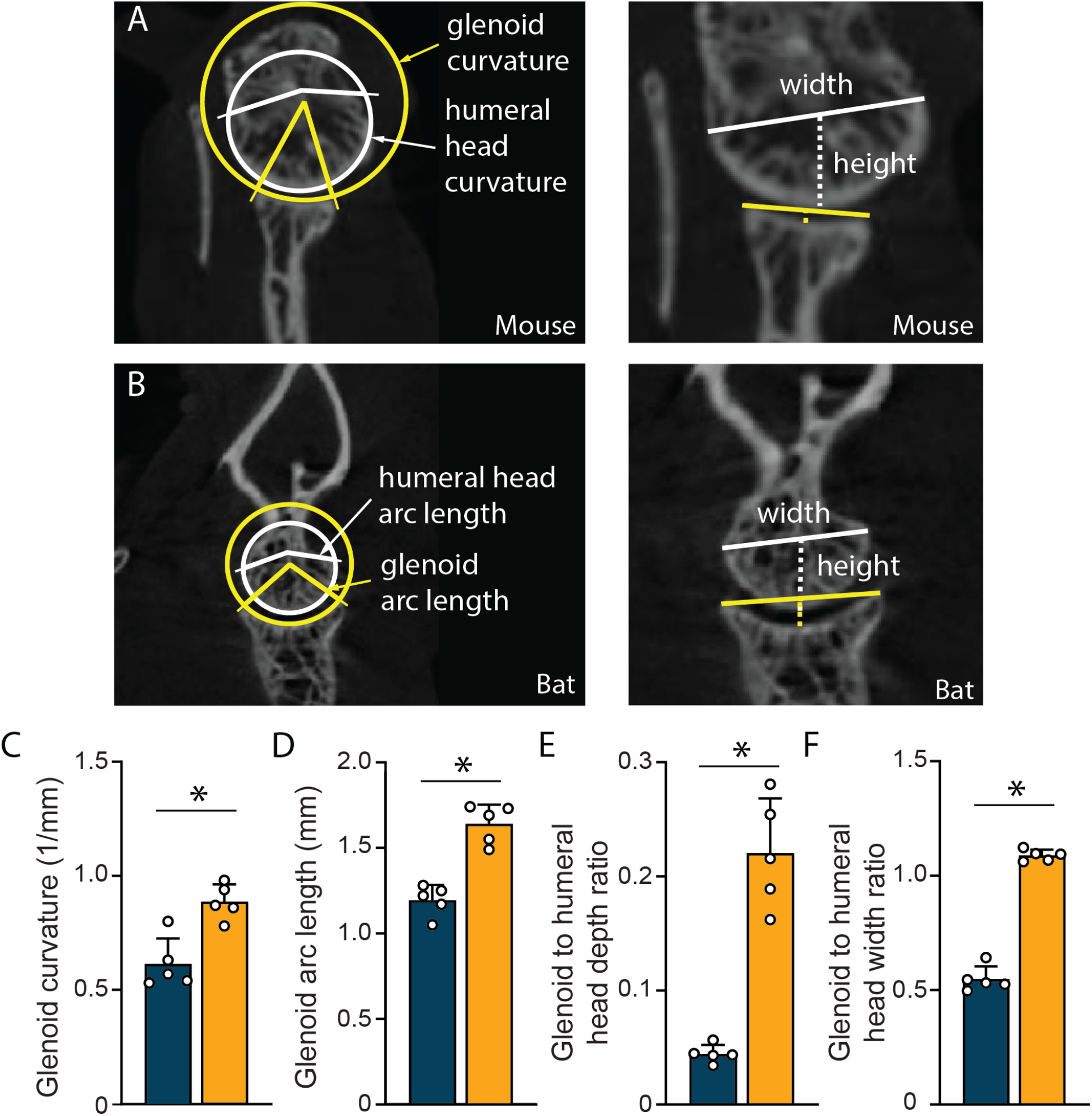
**(A-B)** Representative 3D microCT images of the glenohumeral joint of *C. perspicillata* and mouse showing glenoid curvature and arc length measurements, and glenoid to humeral head depth and width ratios. **(C-D) Glenoid curvature and arc length for mouse and *C. perspicillata***. Glenoid curvature and arc length were significantly higher in bats than in mice. **(E-F) Glenoid to humeral head depth and width ratio**. Both ratios were significantly greater in *C. perspicillata* compared to mice.

### Modeling results predict that the bat shoulder is stable over a dramatically larger range of angles compared to the mouse shoulder

The energy barrier required to overcome shoulder instability was influenced by several factors, including the stiffness of the tendons, peak force, and peak displacement of the humeral head, as well as the depth of the glenoid. The bat shoulder was stable over a dramatically larger range of angles compared to the mouse shoulder (Figure 5). A phase diagram contributes to understanding how this energy barrier for instability varies with tendon insertion angles and glenoid depth (darker shades of red indicate higher energy barriers to instability, Figure 5). The energy needed to dislocate the humeral head of *C. perspicillata* was substantially larger than that in mice, and bat joints were stable for a larger range of shoulder angles. The maximum energy barrier observed for bat supraspinatus and infraspinatus tendon insertion angles was at 60°-70 and 40°-50° respectively (Figure 5 C,D). In the case of mice, the maximum energy barrier for both supraspinatus and infraspinatus tendons was between 30°-40° (Figure 5 A,B). The energy barrier was relatively insensitive to the glenoid angle beyond an instability threshold that is associated with full dislocation of the humeral head from the glenoid; thereafter, the instability is associated with a bifurcation in humeral head position within the glenoid, and the energy barrier for this does not vary with the spatial extent of the glenoid.

**Figure 5.**
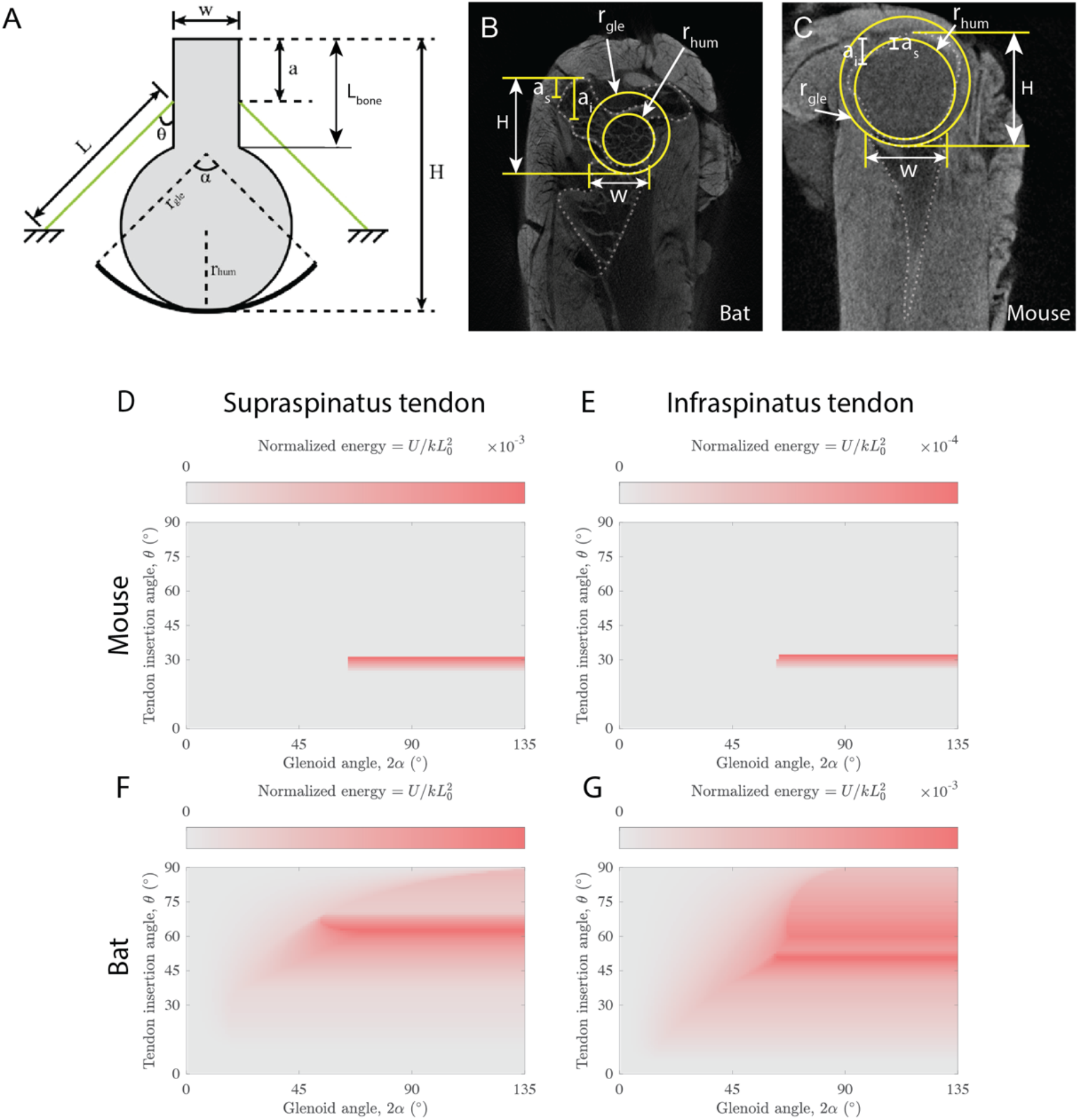
**(A)** Schematic representation of the shoulder instability model. The two-dimensional model considers a humeral head, with a radius *r*_*hum*_ that interfaces with a glenoid with an arc with radius *r*_*gle*_ and angle *α*. **(B-C)** Enhanced micro-CT imaging of *C. perspicillata* and mouse shoulders. Enhanced micro-CT imaging allowed muscle fibers to be visible, making the supraspinatus and infraspinatus attachments easy to visualize. Each of the modeling parameters mentioned above was measured in the transverse plane using IMAGEJ. Each measurement was repeated three times, and an average measurement was taken. **(D-G)** Phase diagram representing the energy barrier that must be overcome for instability of the model shoulder to occur. The energy barrier that resists instability (red) was substantial in bats (F,G) over a larger range than in mice (D,E).

### Bats’ infraspinatus tendons were stronger than those of mice

The cross-sectional area (CSA) of the supraspinatus tendon in bats was significantly larger than in mice. No significant difference in CSA was observed when comparing mouse and bat infraspinatus tendons (Figure 6A). The lengths of the mouse supraspinatus and infraspinatus tendons were approximately the same, whereas in *C. perspicillata*, the infraspinatus tendon was substantially longer than the supraspinatus tendon. Furthermore, the bat infraspinatus tendon was longer than the supraspinatus and infraspinatus tendons of mice (Figure S3B).

**Figure 6.**
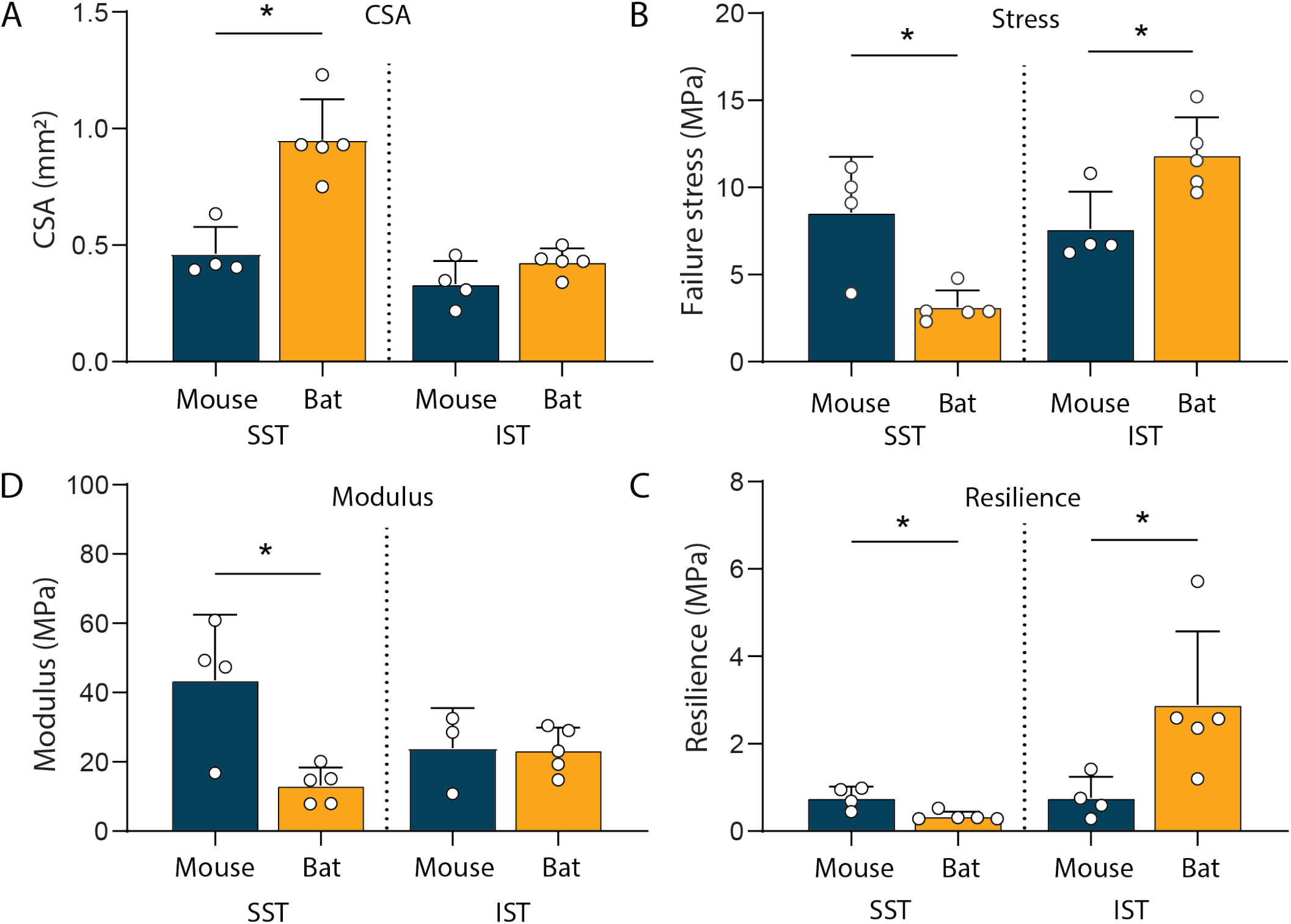
(A-B) CSA measurement in bats and mice. (A) The minimum tendon cross-sectional area for each sample was determined from microCT scans. The cross-sectional area (CSA) of the supraspinatus tendon in *C. perspicillata* was significantly larger compared to that of mice. No significant difference in CSA was observed when comparing the infraspinatus tendons of both species. **Biomechanical testing results**. (B) The *C. perspicillata* supraspinatus tendons exhibited significantly lower failure stress compared to mice, and the *C. perspicillata* infraspinatus tendons showed significantly higher failure stress than that of mice. (C) The modulus of *C. perspicillata* supraspinatus tendons was significantly lower compared to that of mice. There was no significant difference in the modulus of the infraspinatus tendon between *C. perspicillata* and mice. (D) *C. perspicillata* supraspinatus tendons showed significantly lower resilience compared to mice; *C. perspicillata* infraspinatus tendons showed significantly greater resilience compared to mice.

Bat supraspinatus tendons exhibited significantly lower failure stress compared to mice (Figure 6B). In contrast, bat infraspinatus tendons exhibited significantly higher failure stress compared to mice. These results support the idea that the infraspinatus tendons in *C. perspicillata* play a compensatory role, supplementing the function of the supraspinatus tendons. The modulus of bat supraspinatus tendons was also significantly lower compared to mice (Figure 6C). However, there was no significant difference in the moduli of the bat and mouse infraspinatus tendons (Figure 6C). The resilience, or energy absorbed before failure, was significantly lower in bat vs. mice supraspinatus tendons (Figure 6D). In contrast, bat infraspinatus tendons had significantly greater resilience compared to mice (Figure 6D). These results are consistent with a stabilization role for bat infraspinatus tendons during shoulder motion.

The force-displacement curves for both mouse and bat samples had characteristic toe, linear, and yield regions, typical for tendon mechanical behavior under tension (Figure 7 E). Samples failed catastrophically, with little post-yield displacement. Visual examination of the samples showed that the core of the tendon detached from the superficial peritenon tissue as the sample failed. Failure then occurred through bony avulsion, with a small fragment of bone that detached from the humerus remaining on the end of the tendon after failure (Figure 7 A,B). The bony attachments of the supraspinatus and infraspinatus tendons in *C. perspicillata* were qualitatively different from those in mice. Specifically, the greater tuberosity in *C. perspicillata*, where both the supraspinatus and infraspinatus tendons attach, was spherical, with both attachments showing a convex groove shape (Figure 7C). This contrasts with mice, in which the bone was flatter at the tendon attachments, similar to humans (Figure 7D).

**Figure 7.**
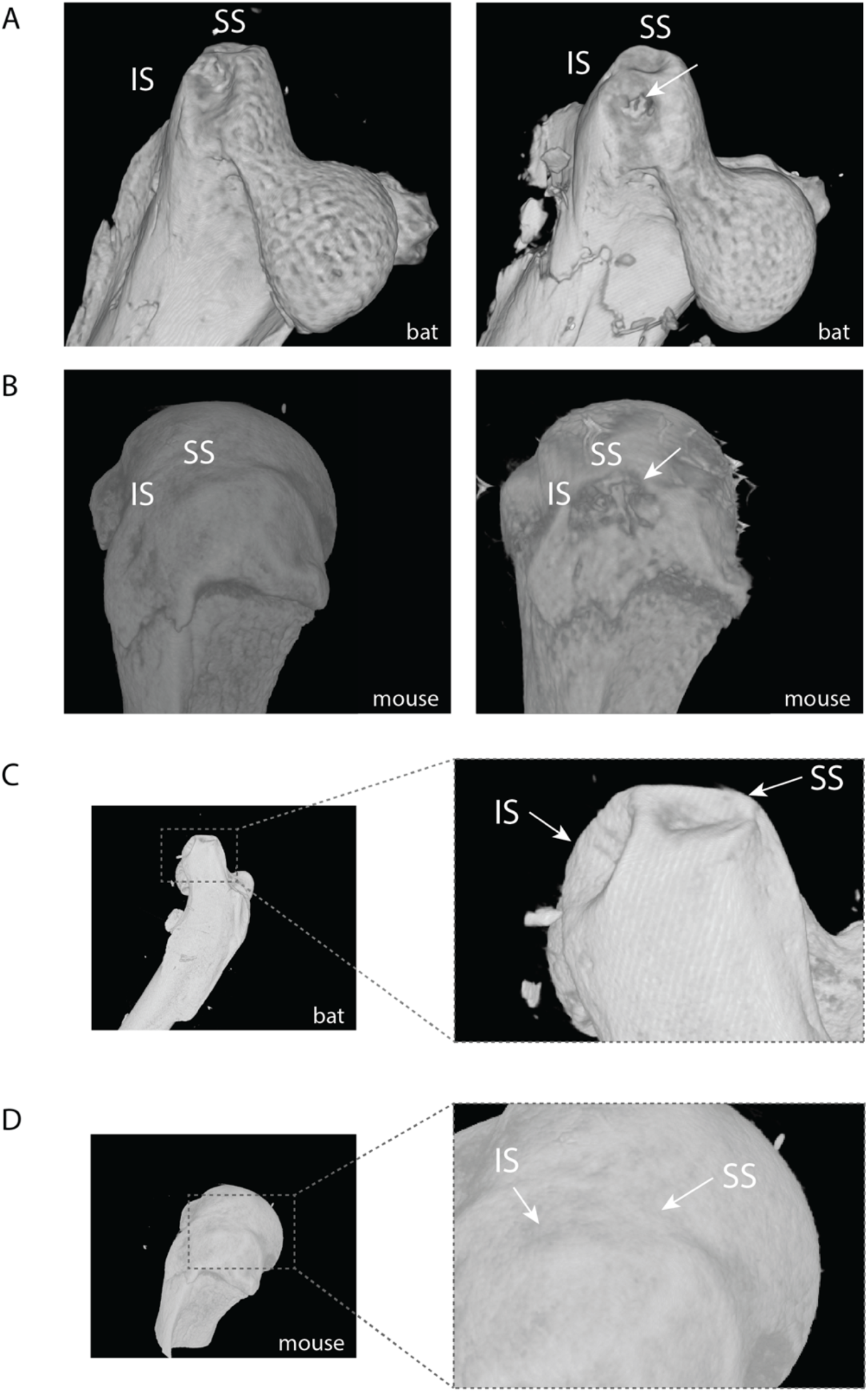
**(A-B)** High-resolution microCT scans and 3D bony reconstructions of humeral head samples from bats and mice post mechanical testing failure showing supraspinatus and infraspinatus attachments. A representative bone avulsion site is visualized (right) for the infraspinatus of bats **(C)** and the supraspinatus of mice (bottom) (**D)**. The mouse attachments are flat, similar to humans, whereas bat attachments have a concave groove shape.

## Discussion

We compared the shoulder anatomy of *C. perspicillata*, typical of that of most bats, to those of quadrupedal mice. During flight, bats regularly employ a wide range of shoulder motions similar to the overhead motions of suspensory primates, and we found unique features of bat anatomy that provide stability to the shoulder. A comparison of anatomy revealed that bat glenohumeral bony structure differs from that of mice in a way that likely reduces stress on the supraspinatus. This is achieved through a larger supraspinatus outlet area and clearance in *C. perspicillata*, suggesting a shift to a greater functional role for the infraspinatus in bats. In addition, we found a unique concavity compression mechanism in *C. perspicillata* that enhances the glenohumeral joint’s stability. The bat’s glenoid has a more pronounced curvature and greater extent, thus encapsulating more of the surface area of the humeral head. The results also imply that the relatively larger infraspinatus in *C. perspicillata* may play a different role than in the mouse.

The scapular index of *C. perspicillata* is greater than those of quadrupedal mice, suggesting a wider, more expansive scapular surface, providing more space for muscle attachments, thus enabling more dynamic movements. A higher infraspinatus index suggests that larger infraspinatus muscles may be required by the bat flight apparatus to contribute to control of external rotation of the humerus at the shoulder without loss of rotational mobility. This is consistent with limited electromyographic data from this muscle in two bat species, *Antrozous pallidus* and *Artibeus jamaicensis* [51].The increased rotator cuff outlet area further emphasizes the importance of this muscle. It is possible that an expanded muscular attachment area might protect against localized stress concentrations by distributing forces generated during overhead motion over a larger surface and decreasing overall stress levels. Alternatively, the relatively smaller supraspinatus muscle we observed in *C. perspicillata* may suggest abduction primarily mediated by this muscle may be less important in bats. The infraspinatus may, indeed, compensate, to some degree, for the small size of the supraspinatus and its limited capacity for abduction [51]. Given that both infraspinatus and supraspinatus stabilize quadrupedal gait, the bat’s enlarged infraspinatus index suggests a larger role for the infraspinatus in the bat’s glenohumeral stability. This trait may also serve to relieve stress on the supraspinatus.

The consistent increase in supraspinatus-acromion clearance during the shoulder’s transition from flexion to extension in both bats and mice indicates a biomechanical advantage conserved across these species. However, the enlarged supraspinatus outlet area we observed in *C. perspicillata*’s may relate to the requirements of extreme wing movements during flight and suggests a potential evolutionary strategy to avoid shoulder impingement syndrome, a pathology seen in overhead-throwing human athletes [52,53]. Impingement syndrome is caused by irritation of the supraspinatus tendon on the undersurface of the acromion [54–56]. Treatment strategies, including surgery, subacromial cortisone injections, or rotator cuff-strengthening exercises, aim to increase the space between the acromion and the rotator cuff tendons [54,55]. In this study, we measured the supraspinatus outlet area and the clearance height but did not examine the outlet area’s shape, which could be useful when considering surgical treatment of shoulder impingement, especially in the case of anterior acromioplasty, which removes bone from the acromion’s undersurface.

The difference between the retroverted glenoid orientation in bats and the anteverted condition in mice is remarkable. Although excessive retroversion is consistently associated with posterior glenohumeral instability and labral tears in humans in some studies [57–61], others indicate that athletes with greater humeral retroversion in their dominant arms experience a reduced incidence of shoulder injuries [62]. A systematic review focusing on overhead-throwing athletes showed a notable increase in humeral retroversion in their dominant vs. non-dominant shoulders [62]. These adaptations occur as a reaction to the stress encountered during overhead throwing activities before the age of 12, a critical period when growth plates are at their peak of activity [62]. Athletes who start participating in these sports post-adolescence do not show this adaptation [62]. The degree of retroversion in bats, analogous to that of high-level overhead-throwing athletes, may represent a balance between enhancing range of motion and stability during extreme overhead motions while minimizing shoulder instability.

The mechanism of concavity compression is crucial for maintaining glenohumeral stability, which, in humans is compromised by traumatic bone loss at the anterior glenoid rim. Currently, defect size and glenoid bone loss are the primary considerations before surgical intervention [20,49]. Although the shape of the glenoid concavity impacts shoulder stability, this factor is not routinely incorporated in clinical assessments of shoulder instability [63,64]. However, recent studies utilizing finite element method (FEM) simulations indicate that glenoid concavity is crucial for glenohumeral joint stability and suggest it be incorporated into clinical use [63,64]. Our results confirmed this concept by showing that the greater glenoid depth of bats corresponds to increased stability compared to mice. This is consistent with characteristics seen in patients with shoulder instability, who often demonstrate a reduction in glenoid concavity and a more flattened glenoid shape[50] [REF]. Hence, we propose that an assessment of individualized concavity might yield a more accurate evaluation of glenohumeral stability for managing bony glenoid defects than the current approach of measuring defect size.

Our instability model builds directly on the concept of concavity-compression and stability. Although it does not capture all anatomical intricacies and degrees of motion of shoulder kinematics, it provides a useful framework for a meaningful comparison between widely different shoulder architectures. It shows, first, that the deep glenoid and larger arc angle of the joint configuration in bats promotes stability over a wider range of tendon insertion angles compared to the mouse. Notably, compared to the mouse, the distinctive anatomy of the bat encompasses a range that includes the maximum energy barrier, indicating that a larger disturbance is required to dislocate the humeral head from the glenohumeral joint. This suggests that the anatomy of bats increases the energy barrier associated with dislocation. We note that the model is idealized and, although it serves as a useful tool for identifying anatomical parameters relevant to human shoulder stability, future studies would benefit from incorporating additional parameters. For instance, instead of using a concentrated force, a future model can incorporate tendons that exert a distributed force over the area at the enthesis footprint and incorporate more complex aspects of glenohumeral joint geometries to expand the range of species modeled.

Mechanical testing of the supraspinatus and infraspinatus tendons from bats and mice showed that the *C. perspcillata’s* infraspinatus tendon is stronger than that of the mouse, supporting our hypothesis that the bat infraspinatus compensates for the smaller, less powerful supraspinatus. A biodiversity-informed approach to shoulder rehabilitation and injury prevention might build on this feature of the bat shoulder and focus on strengthening the human infraspinatus tendon therapeutically. Current treatment approaches typically focus on repairing and strengthening the entire rotator cuff [65]. However, our study suggests that a targeted approach focusing on the infraspinatus could enhance shoulder stability and function, especially when the supraspinatus is compromised. This could lead to more effective and efficient rehabilitation protocols and improved outcomes for individuals with shoulder injuries. Future clinical trials are necessary to test this hypothesis and determine the most effective rehabilitation strategies for shoulder injuries.

The distinctive bony attachments of the supraspinatus and infraspinatus tendons in bats, compared to those of mice, suggest additional aspects of shoulder function that may be implicated in flight mechanics. The indented shape of the attachments, as shown in Figure 7D, might provide a more secure and stable attachment site and a larger area for the tendon/bone interface, which might be beneficial for the repetitive loading by overhead movements associated with numerous wingbeat cycles in long-lived animals. Additionally, the greater tuberosity in bats, where both the supraspinatus and infraspinatus tendons attach, is much larger and protuberant than that in the mouse. This feature likely increases the moment arms of the rotator cuff muscles. In contrast, mice have a flatter attachment and smaller greater tuberosity compared to bats, which is more similar to the human anatomy. Because flatter attachments lead to relatively shorter moment arms for stabilizing rotator cuff muscles, these muscles could be less effective at stabilizing the joint, which would be detrimental during bat flight-generating shoulder motions. To build in this anatomically and functionally distinctive bat proximal humerus, alternative surgical treatment of shoulder injuries in humans could consider modification of the bony geometry of rotator cuff muscle attachment sites.

In summary, this study showed that the glenohumeral joints of bats, the only mammals capable of powered flight, differ substantially from those of a similar-sized quadruped, the laboratory mouse. These findings suggest that bats can serve as a novel model for studying shoulder disease. The traits seen in bats may offer advantages that enhance stability and power during overhead motion. The unique bony geometry of the bat shoulder muscle attachment sites might inspire innovative surgical techniques to improve the treatment of shoulder injuries in humans. Our model studied the effect of attachment geometry on tendon strength, paving the way for future studies that integrate more intricate attachment geometries. Taken together, these findings could ultimately contribute to more effective and efficient rehabilitation strategies and surgical techniques for shoulder injuries in humans.

## Methods

### Sample preparation

All animal procedures were approved by the Columbia and Brown University Institutional Animal Care and Use Committees. Shoulders were harvested from adult (12 weeks) C57BL6/J mice (n=5) and *C. perspicillata* bats (n=5). Note that the average mouse mass was approximately twice that of bats. Glenohumeral joints along with rotator cuff muscles and other surrounding tissues were dissected and then fixed in three different positions: full shoulder extension (“P1”), intermediate posture (“P2”), and full shoulder flexion (“P3”). To consistently identify the angle between the scapular spine and the humerus for each fixation position, quadrupedal (mouse) and flight gait (bat) analyses were used (Figure 1 A and B). Species-specific 3D-printed fixtures were designed to fix samples at each position.

#### MicroCT Imaging

Microcomputed tomography (μCT, Bruker Skyscan 1272) with an energy of 55 kV peaks, an intensity of 145 μA, and a resolution of 19.3 μm was used to scan samples. Each sample at each position was mounted in the microCT scanning sample changer. The data obtained was reconstructed using the provided software (nRecon, Bruker), with alignment optimization and beam-hardening correction applied. The reconstructed image data was visualized using the built-in programs (DataViewer and CTvox, Bruker).

#### Anatomical Measurements

Images were reconstructed with the software (NRecon, Bruker) provided with the CT scanner using alignment optimization and beam-hardening correction. The reconstructed image data was visualized using built-in software (DataViewer and CTvox, Bruker), and bone parameters were measured from reconstructions. The medial end of the scapular spine, superior scapular angle, inferior scapular angle, and center of the glenoid were used as anatomical landmarks. The scapular linear index was determined as the ratio of the width of the scapula (medial aspect of the spine to the glenoid, w) to the length of the scapula (superior to inferior angles, l) (Figure 2A). Supraspinatus and infraspinatus linear indices were calculated in a similar fashion, using the length from the medial aspect of the spine to the superior or inferior angle (Figure 2A). The supraspinatus and infraspinatus area indices were calculated as the area of the respective fossa on the scapula divided by the total area of the scapula in the anteroposterior view, as described previously (Figure 2B) [66]. Glenoid anteversion and retroversion were measured via the anterior facing angle between the body of the scapula and the glenoid face on the axial cross-section. Supraspinatus outlet area was measured as the area posterior to the coracoid, inferior to the acromion, and within the arch of the acromion. The supraspinatus clearance was measured as the vertical distance between the insertion of the supraspinatus on the humerus and the bottom face of the acromion, as seen through the supraspinatus outlet view. Supraspinatus clearance was measured for each sample at each of the three positions. Glenoid curvature was quantified as the inverse of the radius of the best fit circle, measured using the circle tool in ImageJ. The arc length of the glenoid was calculated using the angle from the anterior to the posterior glenoid rim.

#### Mathematical Model of Shoulder Instability

A model of shoulder instability was developed that considered the glenoid shape, the humeral head curvature, and the insertion angles of the rotator cuff tendons. As shown in Figure S1, the two-dimensional model treated the glenoid as an arc defined by a radius *r*_*gle*_ and an *α* angle. The humeral head, with radius *r*_*hum*_, interfaced with a glenoid of width *w*. The total length of the anatomical model, from the glenoid arc to the distal extent of the humeral shaft modeled, was H. Two linearly elastic tendons with unstretched length *L*_0_ and initial length *L* attached to the humeral head at an angle *θ* and a distance *a* from the proximal end of the humerus. We considered the other ends of the tendons to be fixed. Both tendons were modeled as linear elastic with a stiffness *k* when stretched beyond *L*_0_, and exerted no force when not stretched.

To model instability through destabilizing forces, a concentrated force 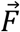 was applied in a quasistatic manner (Figure S1). The force was increased linearly until the humeral head became unstable, at which point the force needed to displace it further in the glenoid decreased. To maintain the system in a static state, three other forces were present: the tension from the tendons, 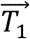 and 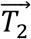, and the normal force 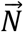 exerted by the static glenoid. As a result of 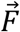 the humeral head slides along the glenoid (its component in the x-axis marked as δ at an angle *γ*. As the magnitude of 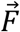 increased, the angle of humeral head relative to the glenoid also increased, until a closed-form solution for the four balanced forces could no longer be found, indicating shoulder instability.

In a system with specified geometry (i.e., *r*_*gle*_, *r*_*hum*_, *w, H, a, L*_0_, *L, k, α*, and *θ* are defined), the correlation between the displacement of the bottom end of the humeral head, δ, and the magnitude of the external force, 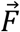, can be determined through the following steps. We defined *β* as the angular displacement of the tangent point between the humerus and the glenoid, *γ* as the tilting angle of the humerus-glenoid assembly, and 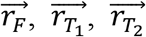 as the displacement vectors from the tangent point to the points where 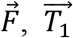 and 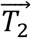 are exerted, respectively. For a specific value of *β*, 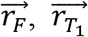 and 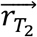 are all functions of *γ*, as are the 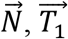 and 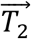. The balance of forces and moments with respect to the tangent point on the humeral head - glenoid assembly are as follows:

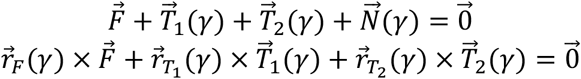

Solving the above system of vector equations for 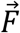 and *γ* yields the necessary magnitude of 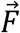 to stabilize the system for a specific bottom angular displacement *β*. Shoulder instability was identified when the calculated *γ* exceeded 90°, when the computed force was non-positive, or when no closed-form solution could be determined for 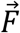. Hence, for a system with specified geometry, the correlation between δ and 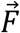 can be derived. Similarly, the relationship between δ and *γ* can also be derived.

With *F*(δ)represented as a function of δ, the correlation between δ and the energy, *E*, stored in the system for a known system geometry can be calculated using the following equation:

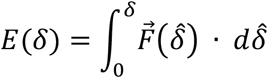

where force 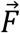is exerted and is a function of δ. The maximum energy that a system can store is noted as *E*_*max*_. *E*_*max*_ can be calculated for any system, where the model parameters can be adjusted to compare system stability. We compared 4 scenarios of shoulder instability: (1) mouse supraspinatus tendon, (2) mouse infraspinatus tendon, (3) bat supraspinatus tendon, and (4) bat infraspinatus tendon (Table S1). A phase diagram, where a darker region represents higher *E*_*max*_ was constructed, with varying *θ* as the vertical axis and *α* as the horizontal axis. The geometrical parameters were altered for all four scenarios and different phase diagrams were compared. The parameters used for all simulations are shown in Table S1 and were measured using reconstructed microCT images of mouse and bat scans as shown in Figure S2. The stiffness of each tendon was determined through mechanical testing, as in the mechanical testing section below.

### Mechanical testing

#### Sample preparation

All animal procedures were approved by the Columbia University and Brown University Institutional Animal Care and Use Committees. Supraspinatus tendon-to-bone attachment units and infraspinatus tendon-to-bone attachment units (humerus-tendons-muscle) were harvested from adult (>10 weeks) C57BL6/J mice (n=5) and *C. perspicillata* bats (n=5). After dissection, samples were wrapped in PBS and fresh-frozen and stored at –20°C.

#### Tendon cross-sectional area measurement using micro computed tomography (microCT) imaging

To prepare for scanning, the distal end of the supraspinatus-humerus unit was embedded in 2% agarose (Sigma-Aldrich) and mounted in the microCT scanning sample changer. This ensured that the tendon enthesis specimens were suspended freely and aligned with the scanning axis. MicroCT scans were conducted (μCT, Bruker Skyscan 1272) with an energy of 55 kV peaks, an intensity of 145 μA, and a resolution of 19.3 μm. The data obtained was reconstructed using the provided software (nRecon, Bruker), with alignment optimization and beam-hardening correction applied. The reconstructed image data was visualized using the built-in programs (DataViewer and CTvox, Bruker).

The minimum tendon cross-sectional area for each sample were determined from microCT scans. These scans were conducted on samples before mechanical testing and analyzed using built-in image processing algorithms (CTAn, Bruker). A thin sheet of kimwipe paper was placed between the supraspinatus and infraspinatus tendons to differentiate them post-scan (Figure 3.2 A). The minimum cross-sectional tendon area was identified by thresholding transverse slices of the tendon, calculating the area covering the tendon, and choosing the smallest area that is within 500 µm of the tendon insertion site (Figure 3.2 A).

#### Tensile testing

Samples underwent mechanical testing in a saline bath at 25°C using a table-top tensile tester (Electroforce 3230, TA Instruments) equipped with a 10 lb load cell (TA Instruments). Prior to testing, the supraspinatus muscle was carefully scrapped off from the supraspinatus-humerus unit. These samples were secured using custom 3D-printed fixtures [67] specifically designed for bat shoulder anatomy (Figure 3.1 A). To ensure a firm grip, the tendon was sandwiched between two layers of thin Kimwipe paper and adhered using a drop of cyanoacrylate adhesive (Loctite, Ultra Gel Control) before mounting to the custom grips. Samples were secured in fixtures and tested in an orientation corresponding to 90° shoulder abduction. The testing regimen involved preconditioning the samples with a sinusoidal load ranging from 0.05 to 0.2 N for 5 cycles, followed by a 2-minute resting period. Thereafter, samples were strained until failure at a rate of 0.1% per second. Each specimen consisted of both the supraspinatus and infraspinatus tendons attached to the humerus. The supraspinatus tendons were tested first, after which the infraspinatus tendons were tested using the same mounting methods.

#### Biomechanical analysis

Enthesis structural properties, such as failure load (referred as “strength” in the text), stiffness, and toughness (i.e., the area under the curve through failure load) were determined from load-deformation curves. Stiffness was determined from the force-elongation curve using random sample correlation (RANSAC). Data was first trimmed to remove data below 10% and above 95% of maximum load to identify the region of interest. Then, two points were selected at random and a line was drawn between them for n = 1000 iterations. All data points within a threshold range of 0.5% of the robust fit stress at the 80th percentile were considered as within an acceptable range of the best fit line. Of the n iterations, the iteration with the most inliers was deemed the best fit. This approach represents a “robust” fit, which compared to a least squared errors fit, minimizes the effect of outlier points on the best-fit line. The best fit was confirmed by visual inspection for each force-elongation curve. The “yield” point was determined as the point where the smoothed data first deviated from the RANSAC fit line by 0.1% of the expected load. To confirm appropriate identification, the yield point was also determined manually (i.e., the user manually selected the “yield” point on the graph). The manual selection was repeated 10 times to account for user error. Energy was calculated as the area under the load deformation curve up to the yield point.

### Statistical analysis

Statistical analysis was performed for all experiments using unpaired t-tests with Welch’s correction (GraphPad Prism 7). All data are shown as mean ± standard deviation. The threshold for statistical significance was defined at p < 0.05. Sample size, n, represents biological replicates.

## Acknowledgments

This work was supported by National Institutes of Health (R01 AR080717 and R01 AR077793) and Air Force Office of Scientific Research (FA9550-12-1-0301 DEF) and a Brown University Seed Award to SMS

## Author Contributions

Conception: I.K., S.T., G.M.G., S.M.S.;

Study design: I.K., S.T. G.M.G., S.M.S.;

Performed experiments/modeling: I.K., J.K., S.L.,

Critical interpretation of results: all authors; Drafted the manuscript: I.K., S.T.;

Read and edited the manuscript: all authors.

