## Supplementary material for "Musculoskeletal architecture of the shoulder: A comparative anatomy study in bats and mice informing human rotator cuff function": Supplmental Data

**Table of contents**

Supplementary Note 1: Contrast enhanced microCT imaging

Fig. S1. Schematic representation of the shoulder instability model

Fig. S2. Enhanced micro-CT imaging of mouse and bat shoulders

Fig. S3. Mechanical testing workflow

Fig. S4. CSA measurement and representative strength (force)-displacement curve

Table. S1. Shoulder instability model parameters

**Supplementary notes**

Contrast enhanced microCT imaging

Bat and mouse shoulder joints (including the humerus, scapula, surrounding rotator cuff muscles and tendons) were dissected and fixed in 4% PFA before undergoing PTA staining and micro-CT imaging. A PTA solution was prepared by dissolving approximately 5 ml of phosphotungstic acid hydrate, a dry powder (Alfa Aesar), in 200 ml of 70% ethanol, ensuring that the PTA powder dissolved completely in ethanol. Placing the samples directly from buffered formalin storage into alcohol-based PTA can cause tissue cracking due to rapid dehydration. Therefore, the samples were serially dehydrated in ethanol, starting from 30%, progressing to 50%, and then to 70%, with each dehydration step lasting overnight. After serial dehydration, individual samples are submerged in a few milliliters of the PTA solution in small, sealed, flat-bottomed plastic vials and left in the stain for 1-7 days. A quick scan was conducted each day to assess the penetration depth of the stain. After ensuring that the stain has penetrated through and the soft tissues are visible in the scan, the samples are removed from the PTA solution, placed on paper tissue, and superficially dried for a few seconds. They are then wrapped in thin strips of soft paper tissue and prepared for micro-CT imaging. Once the paper-wrapped samples were positioned in the plastic tube, a few drops of tap water were added to moisten the paper surrounding the sample, preventing it from drying during the scanning process. The same micro-CT imaging protocol mentioned above was followed to reconstruct the scans. The enhanced micro-CT imaging allows muscle fibers to be visible, making the supraspinatus and infraspinatus attachments easy to visualize. Each of the modeling parameters mentioned above was measured in the transverse plane shown in Figure S2 using ImageJ software. Each measurement was repeated three times, and an average measurement was taken.

**Supplementary Figures**

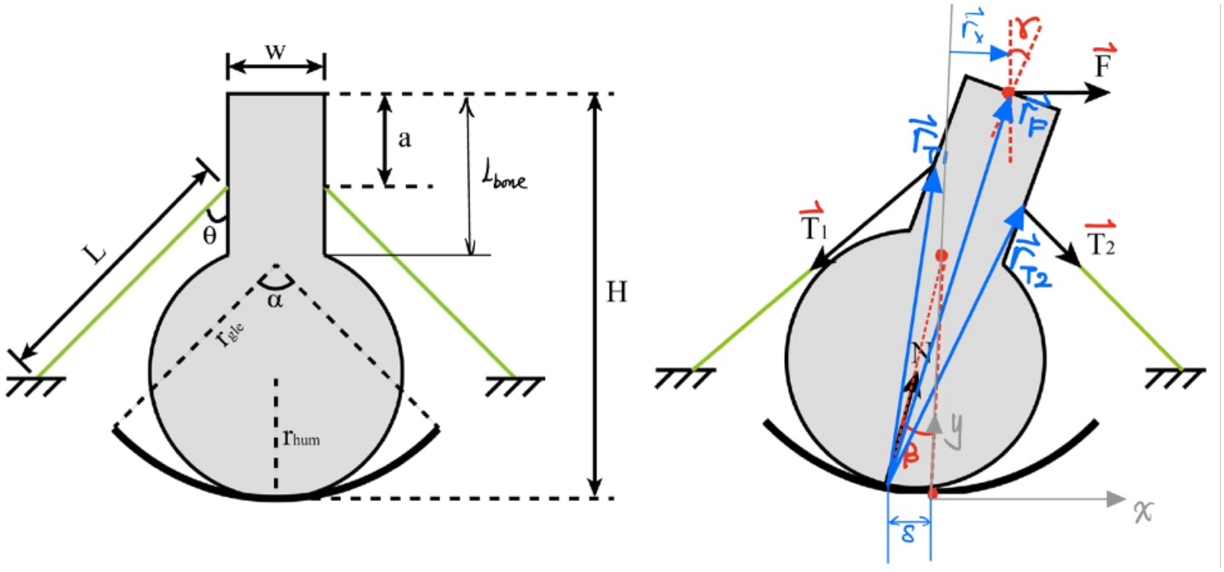


**Figure S1.** Schematic representation of the shoulder instability model. The two-dimensional model considers a humeral head, with a radius $rhum$ that interfaces with a glenoid with an arc with radius $rgle$ and angle $\alpha$. A concentrated force $\vec{F}$ was applied in a quasistatic manner and increased linearly until the humeral head became unstable.

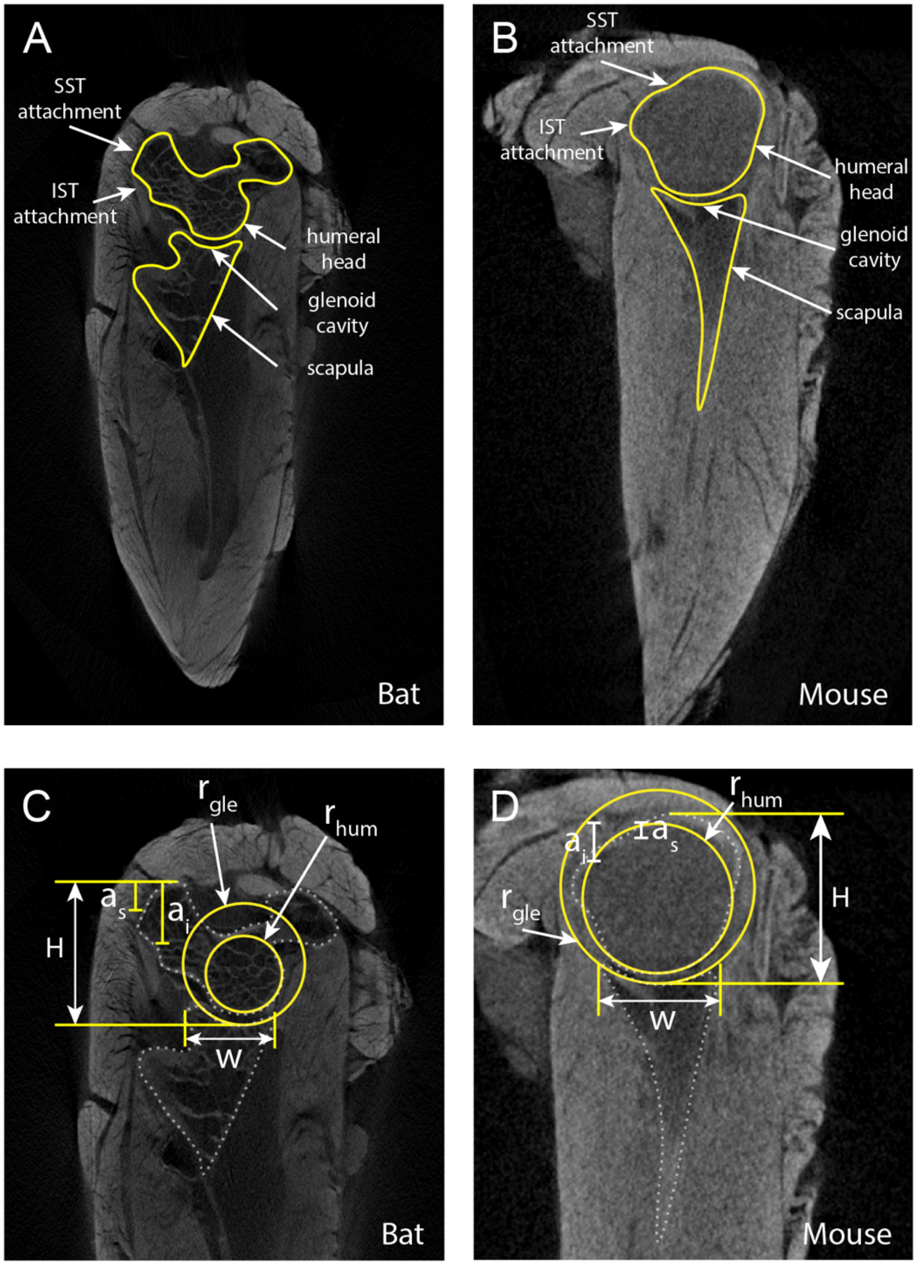


**Figure S2.** Enhanced micro-CT imaging of mouse and bat shoulders. Enhanced micro-CT imaging allowed muscle fibers to be visible, making the supraspinatus and infraspinatus attachments easy to visualize **(A,B)**. Each of the modeling parameters mentioned above was measured in the transverse plane using IMAGEJ. Each measurement was repeated three times, and an average measurement was taken **(C,D)**.

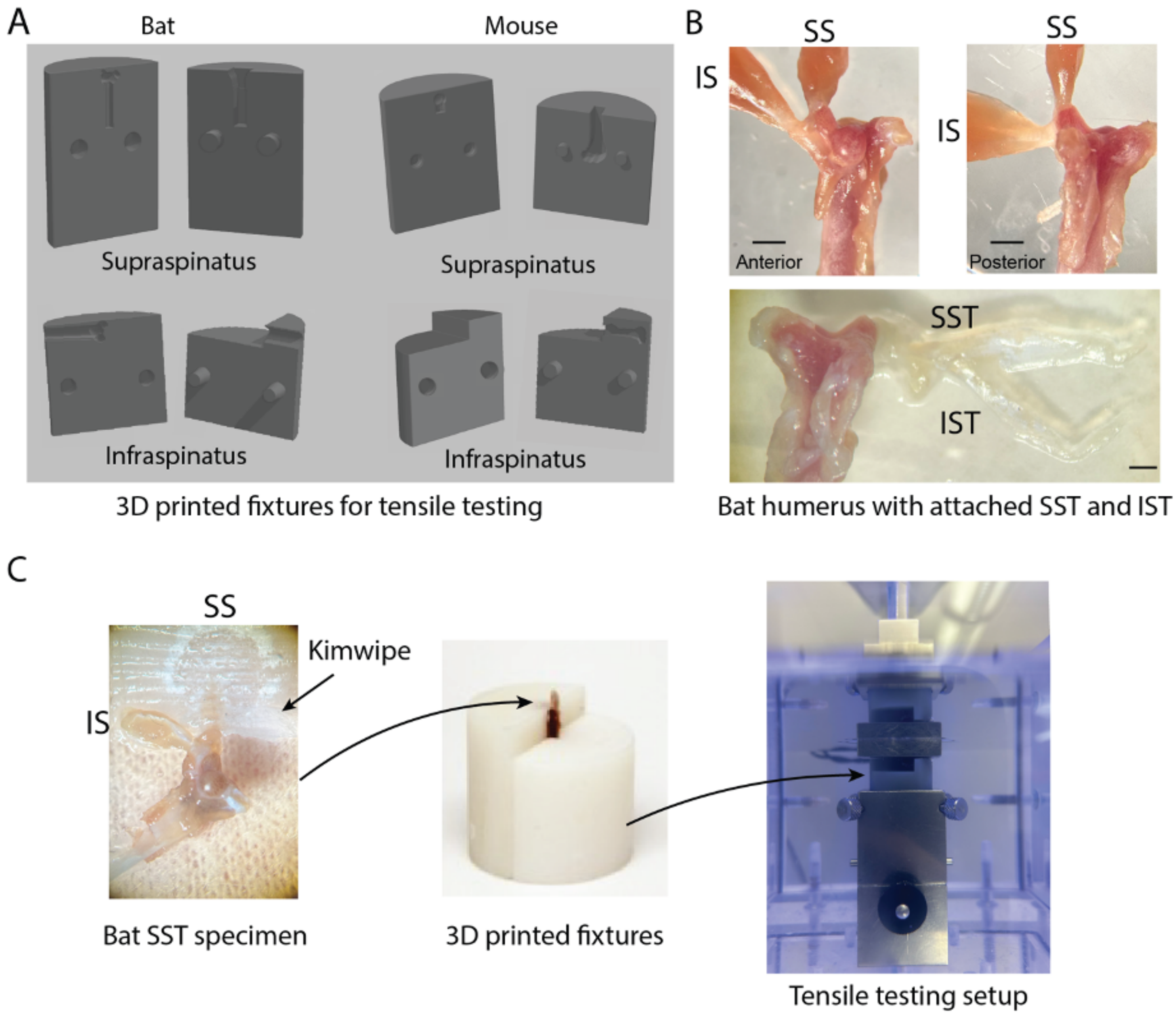


**Figure S3. (A)** Custom 3D-printed fixtures were made to test the supraspinatus and infraspinatus tendons of bats and mice. **(B)** Anatomy of the bat showing humerus with supraspinatus and infraspinatus tendons attached. (Scale: 1mm). **(C)** Prior to testing, the supraspinatus muscle was carefully scrapped off from the supraspinatus-humerus unit, and the tendon was sandwiched between two layers of thin Kimwipe paper and adhered using a drop of cyanoacrylate adhesive before mounting to the custom grips. The same workflow was followed for the infraspinatus tendons.

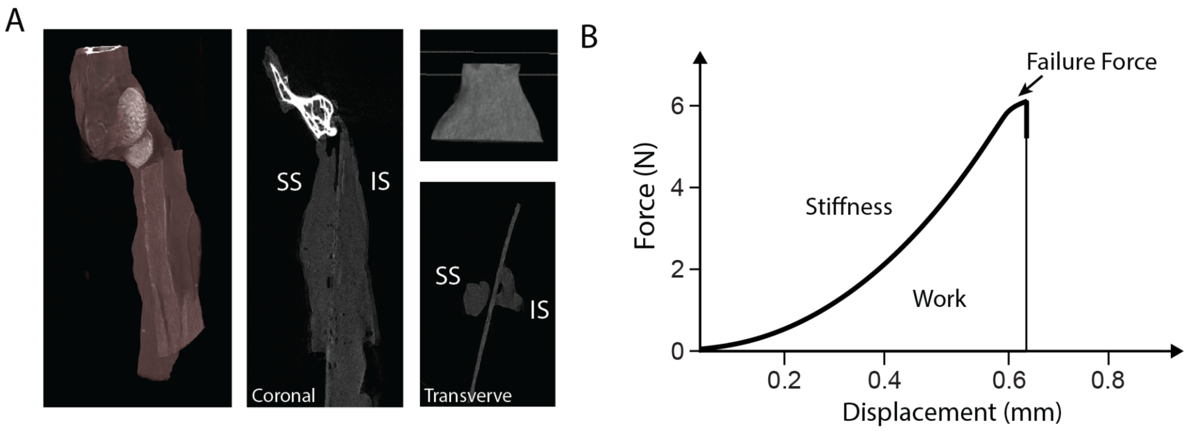


**Figure S4. (A)** The cross-sectional area (CSA) of the supraspinatus tendon in bats was found to be significantly larger compared to that of mice. No significant difference in CSA was observed when comparing the infraspinatus tendons of both species. **(B)** Representative strength (force)-displacement curve.

| Parameters | Bat SST | Bat IST | Mouse SST | Mouse IST |
|---|---|---|---|---|
| $r_{hum}$ (mm) | 0.84 | 0.84 | 1.00 | 1.00 |
| $r_{gle}$ (mm) | 1.14 | 1.14 | 1.67 | 1.67 |
| $w$ (µm) | 1,060.33 | 1,060.33 | 1,591.33 | 1,591.33 |
| $H$ (µm) | 2,804 | 2,804 | 1,972 | 1,972 |
| $a$ (µm) | 572 | 1,248 | 239.33 | 530.33 |
| $L$: Length at 3% strain (mm) | 10.57 | 12.36 | 7.21 | 5.15 |
| $L_0$: Length unstretched (mm) | 10.26 | 12 | 7 | 5 |
| $k$: stiffness (N/mm) | 5.41 | 5.47 | 11.00 | 2.18 |

**Table S1.** Shoulder instability model parameters.
